# Brief communication - *Phaeodactylum tricornutum* is a bona fide calcifying diatom

**DOI:** 10.64898/2026.10.12.758572

**Authors:** George Blundell-Hunter, Lucie Paoli, Oscar Marsden, Elena Kazamia, Philippe Mozzanega, Aranza Diaz-Ramos, Nanakow Baiden, Emily McGibbon, Ekta Kumari, Ruby Samuels, Jonathan Collard, John Waite, Ian Hu

## Abstract

*Phaeodactylum tricornutum* is an unusual member of the clade of diatoms, eukaryotic marine brown algae that possess a distinct cell wall made of silica. *Phaeodactylum* is a rare exception in diatom biology in that it does not require silica throughout its whole life cycle. Here, we demonstrate that *Phaeodactylum* has a calcifying activity, which is highly unusual in unicellular eukaryotes.

## Introduction

Diatoms are a monophylyl group of marine photosynthetic eukaryotes. They stand out with their tertiary lineage of their chloroplasts, their complex and unorthodox multi-layered endomembrane systems, their photosynthesis of a significant portion of total global primary oxygen, and for their unique silica-based cell wall and biology. *Phaeodactylum tricornutum* is a commonly used model organism for eukaryotic algae for its genetic tractability. Being the black horse of the class, *Phaeodactylum* is an exception to the obligate silica requirement of the diatoms throughout its life cycle. We report here a biological activity, calcification, that has not been reported in unicellular eukaryotes outside the clade of coccolithophores - a genetically unrelated class of Haptophyte phytoplanktons - whose calcified outer shell deposition forms geological phenomena such as the White Cliffs of Dover.

## Results

### *Phaeodactylum* generates insoluble deposit when grown in closed polycarbonate vessels, polycarbonate surfaces in open ponds, and standard polypropylene tissue culture flasks

We routinely grow Phaeodactylum cells between 150 mL and 1000 mL in volume in our bubbling polycarbonate GL 45 system photobioreactor (Fig. 1a) (Hu et al., in prep) to reduce the influence of silica from glass surface [1]. Curiously, when the cells were cultured in less nutrient-replete media such as f/2 or L1* [2], we saw translucent white deposit formation at the meniscus of the media and air, and where air bubbles would break (Fig. 1b), a phenomenon that is not present in mock bottles without Phaeodactylum cells. Initially we suspected that this was a metabolic product due to the genetic modifications we performed, but we saw identical phenomena on our wild type controls as well. We attempted to clean up this material by washing with first saline, then purified water, then hexane and ethanol. The material has very poor solubility in both organic solvents, though when flushed with heated ethanol, the deposit could be removed from the vessel. The product was not dissolved though, as after transferring to a regular 1.7mL microcentrifuge tube, the deposit would very quickly precipitate to the bottom of the ethanol layer (Fig. 1c). We observed that the deposit had exceptionally high solubility when washed with dilute hydrochloric acid (not shown).

**Fig. 1.**
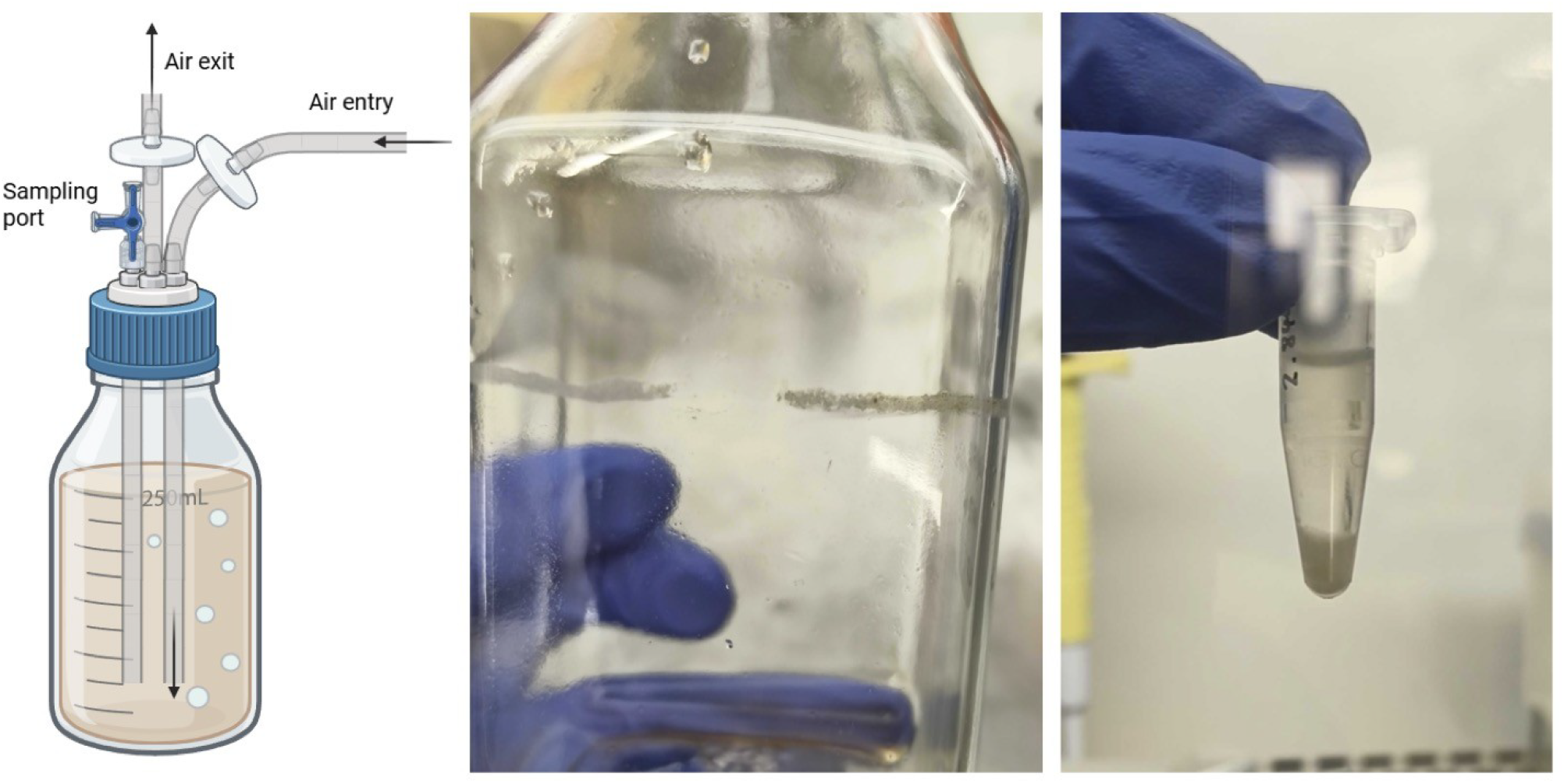
Deposit formation in 250mL-scale bottle bubbling photobioreactor. Left, 1a, schematic of the bottle bubbling photovioreactor. Middle, 1b, the deposit attached to the bottle under white light. Right, 1c, the deposit after being moved into a microfuge tube through ethanol.

We further tested growth of Phaeodactylum in open ponds 5L and 150L in size. The 5L pond is wholly constructed in polycarbonate, whereas the shell of 150L ponds were fabricated in fibreglass with polycarbonate paddle wheels. For inoculating the open ponds, cells were routinely grown in 1/20 of the target volume of the pond in a highly nutrient-replete media that we call Artificial Sea Water Media (ASWm) [3] up to early stationary phase in appropriate vessels - a bubbling photobioreactor or a hanging bag - and then the whole volume of the inoculum would be transferred into the pond with media premade inside. We made the same observation in the ponds, that is when the cells were grown in either f/2 or L1*, a similar insoluble translucent deposit would form on the polycarbonate surface in significant contact with air. In the 5L ponds’ case this was on the inner pond wall the whole way above and below the menisci, and in 150L it presented on the paddle wheel (Figs. 2a - 2c).

**Fig. 2.**
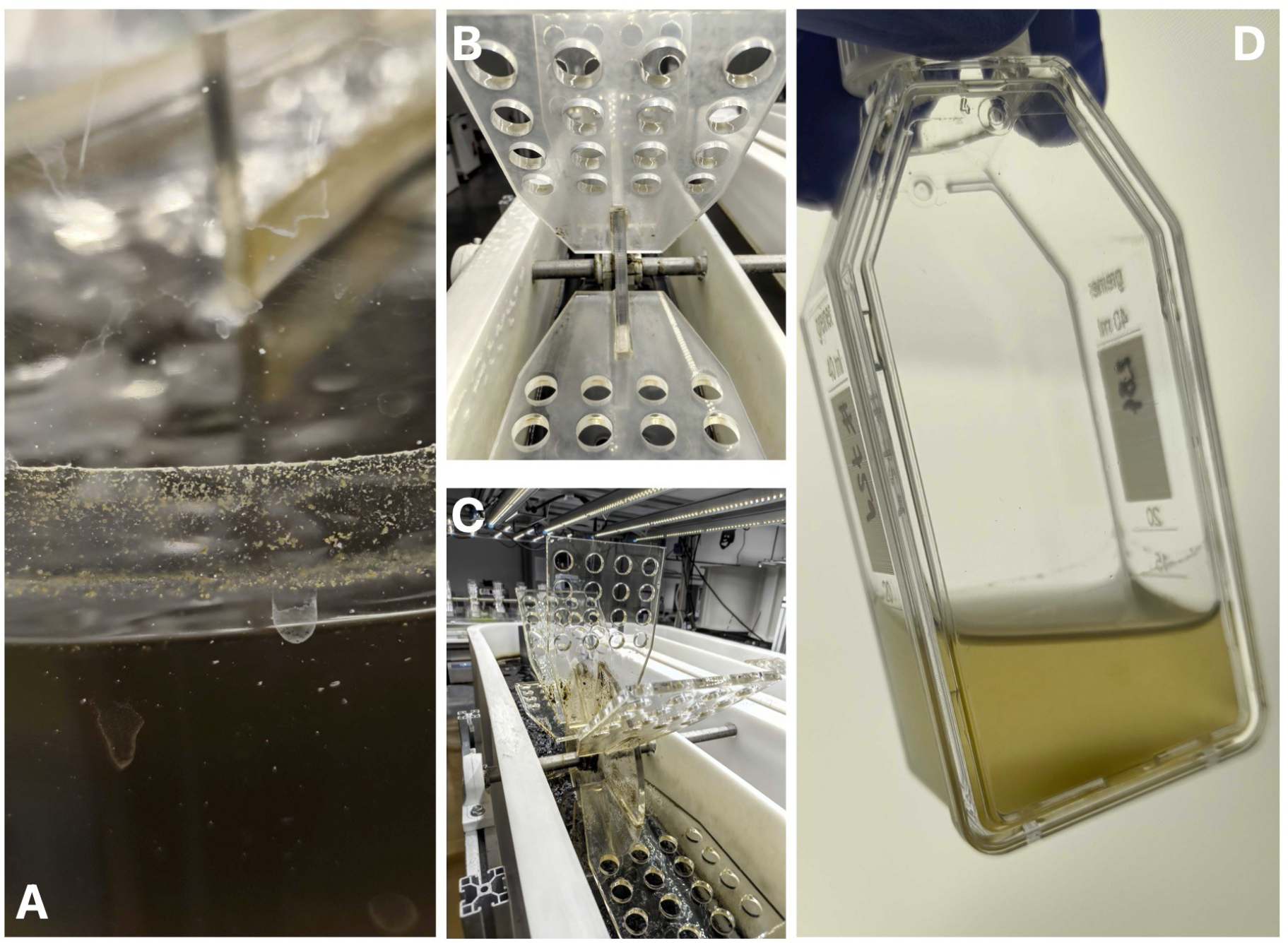
Deposit formation in other plastic vessels. 2a, deposit formation in 5L raceway pond. 2b, deposit formation on the paddle wheel of a 150 L raceway pond. 2c, paddle wheel without deposit for reference. 2d, deposit formation on a T25 tissue culture flask.

As we began suspecting that deposit formation may have relation to media composition, we then performed cell culture with f/2 in regular tissue culture flasks, T25 for suspension cells, under common laboratory conditions: 50 mmol/m2 illumination on a 16/8 light/dark cycle at 18°C, gentle orbital shaking. The T25, being of polypropylene rather than polycarbonate, reproduced the similar phenomenon of white deposition at the media-air menisci (Fig. 2d). We note that in all the experiments, regardless of scale, once the media is autoclaved, we invariably observed salt precipitation in the autoclaved vessel, and this media would no longer generate the deposit.

### Chemical analyses of the deposit

We dried and harvested the deposit from the 150L pond paddle wheels, then performed an exhaustive organic solvent test, and failed to dissolve the material in methanol, ethanol, isopropanol, hexane, heptane, isooctane, mineral oil, diethyl ether, methyl tert-butyl ether, dichloromethane, chloroform, dimethylforamide, dimethylsulfoxide, acetone, and acetonitrile. We analysed the material with ICP-MS (excluding C, H, N, O). The material is disproportionally high in calcium and magnesium atoms (Table 1) considering the media was analogous to a fertilised seawater. We then performed a higher weight dissolution of the material in 0.1N HCl, and to our surprise we saw gas bubble formation (not shown), providing strong evidence that this is a carbonate divalent salt solubilising event, with the counter ion being a combination of calcium and magnesium.

**Table 1.**
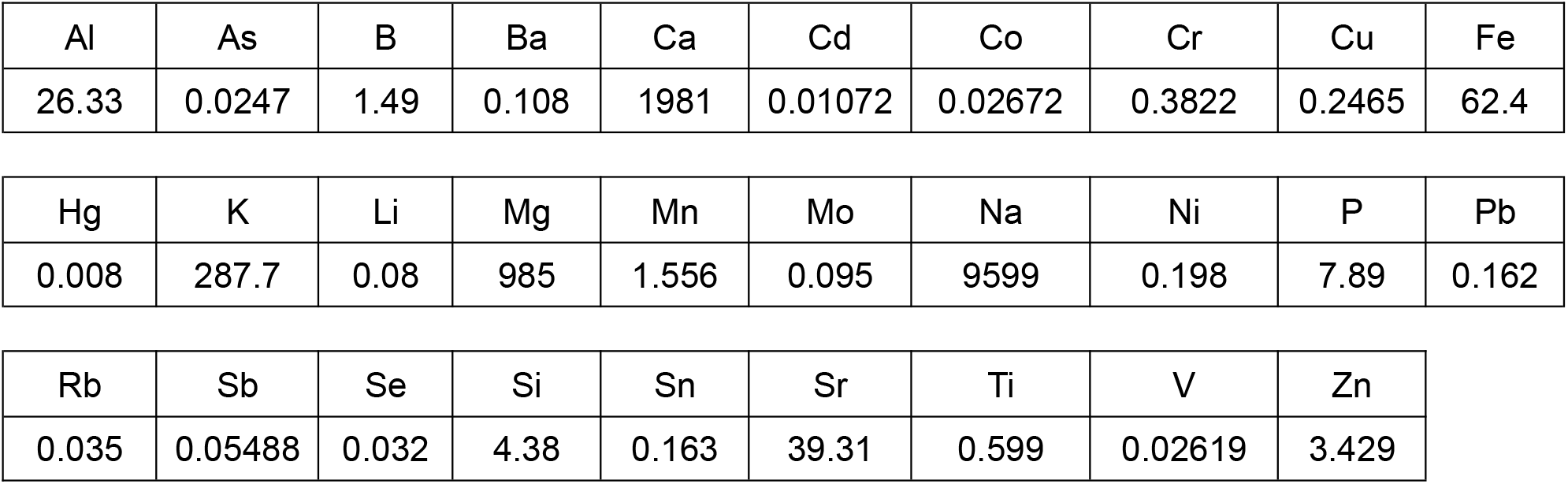
ICP-MS results of the deposit. Unit is mg/L. Sample preparation is described in the Methods section.

|  |  |  |  |  |  |  |  |  |  |
| --- | --- | --- | --- | --- | --- | --- | --- | --- | --- |
| Al | As | B | Ba | Ca | Cd | Co | Cr | Cu | Fe |
| 26.33 | 0.0247 | 1.49 | 0.108 | 1981 | 0.01072 | 0.02672 | 0.3822 | 0.2465 | 62.4 |
| Hg | K | Li | Mg | Mn | Mo | Na | Ni | P | Pb |
| 0.008 | 287.7 | 0.08 | 985 | 1.556 | 0.095 | 9599 | 0.198 | 7.89 | 0.162 |

|  |  |  |  |  |  |  |  |  |
| --- | --- | --- | --- | --- | --- | --- | --- | --- |
| Rb | Sb | Se | Si | Sn | Sr | Ti | V | Zn |
| 0.035 | 0.05488 | 0.032 | 4.38 | 0.163 | 39.31 | 0.599 | 0.02619 | 3.429 |

### Calcite-like crystalline cell clumps observed under microscopy

To further understand the nature of the deposit, we used a hand rotary tool to remove pieces from a 250mL polycarbonate bottle with the deposit in question and observed it with a high-power stereomicroscope (VHX-X1, Keyence Ltd) (Fig. 3). To our surprise, the deposit is composed of single Phaeodactylum cells and cell clumps that seemingly are fused through means yet unknown, presenting with a conglomerate phenotype where the distinction between individual cells could no longer be distinguished completely.

**Fig. 3.**
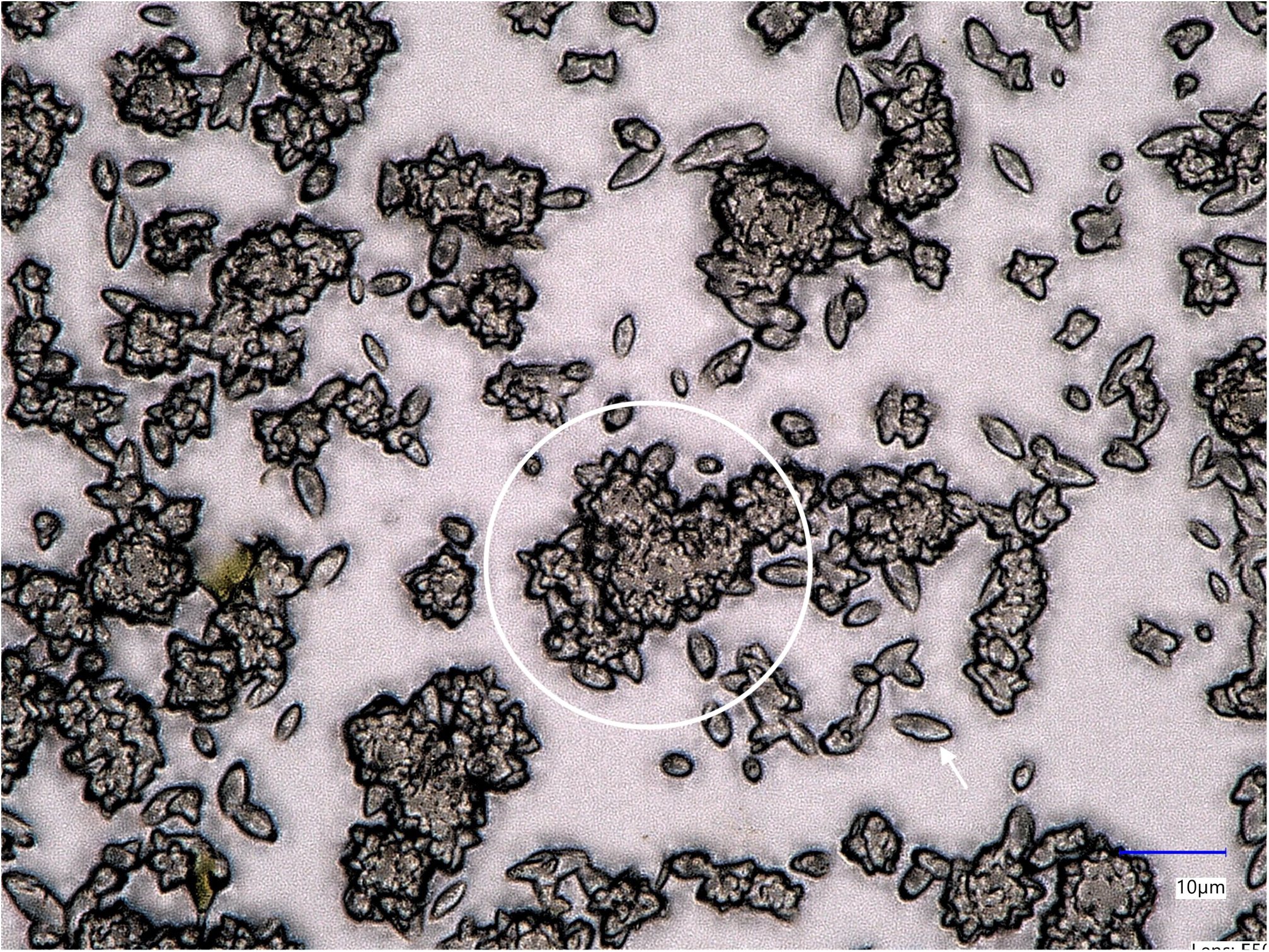
calcification-like deposit under stereomicroscopy. White arrow, example of a single cell. White circle, example of a cell clump showing opaque extracellular material binding individual cells together.

### Initiation of calcification seems to be determined by metabolic state as well as a diffusible chemical signal

To further interrogate the trigger to this highly unusual behaviour, we made a series of outgrowth, spiking, and media replacement experiments with the formation of deposit as output. The results are summarised in Table 2. We observe that a culture can only generate a fixed amount of deposit, which seems to be in proportion to the initial nutrient content of the culture that eventually goes into stationary phase as well as the final OD, but not necessarily in proportion to cell number. When two cultures were inoculated with the same amount of inoculum, the one with higher N/P, if hit stationary phase, would see higher OD and more deposit, though cell counts would be roughly the same. Once a culture produced the deposit and stopped, the production could not be reinitiated in the same media by addition of extra nutrients.

**Table 2.** Summary of Phaeodactylum outgrowth, spiking, and media replacement experiments related to carbonate deposit formation.

| Type | Treatment | Outcome |
| --- | --- | --- |
| Outgrowth | f/2 | Deposit formation at early stationary phase above certain threshold of inoculation density |
| Outgrowth | L1* | Stronger deposit formation at early stationary phase above certain threshold of inoculation density |
| Outgrowth | ASWm | No deposit formation |
| Spiking (stationary phase) | Mock | Culture sustenance, no deposit formation |
| Spiking (stationary phase) | f/2 | Culture crash |
| Media replacement | Seawater | Culture crash |
| Media replacement | f/2 | Healthy culture, deposit formation |
| Media replacement | L1* | Healthy culture, stronger deposit formation |
| Media replacement | ASWm | Hyper-dense culture, no deposit formation |

## Discussion

### Metabolic trigger of the calcifying event

We initially considered that the trigger is the metabolic switch into stationary growth phase. However the media replacement result suggests that the trigger may be much closer to the nutrient starvation trigger itself, as when a stationary phase culture is washed in seawater and replaced in an equal volume fresh low nutrient media, the phenomenon could be triggered again without cells reentering log phase. We posit then that the trigger is nutrient depletion sensed by the cells, with a negative feedback inhibitor in the form of an extracellular small molecule diffusible signal that is secreted by the cells when they are forced to enter stationary phase by nutrient depletion (Fig. 4). Murphy et al. [4] has already demonstrated that glutamate acts as an extracellular signaling molecule in Phaeodactylum, providing strong evidence that mechanistically monocellular diatoms are able to use small molecules for intercellular communications, a mode generally regarded as exclusive to larger, multicellular organisms.

**Fig. 4.**
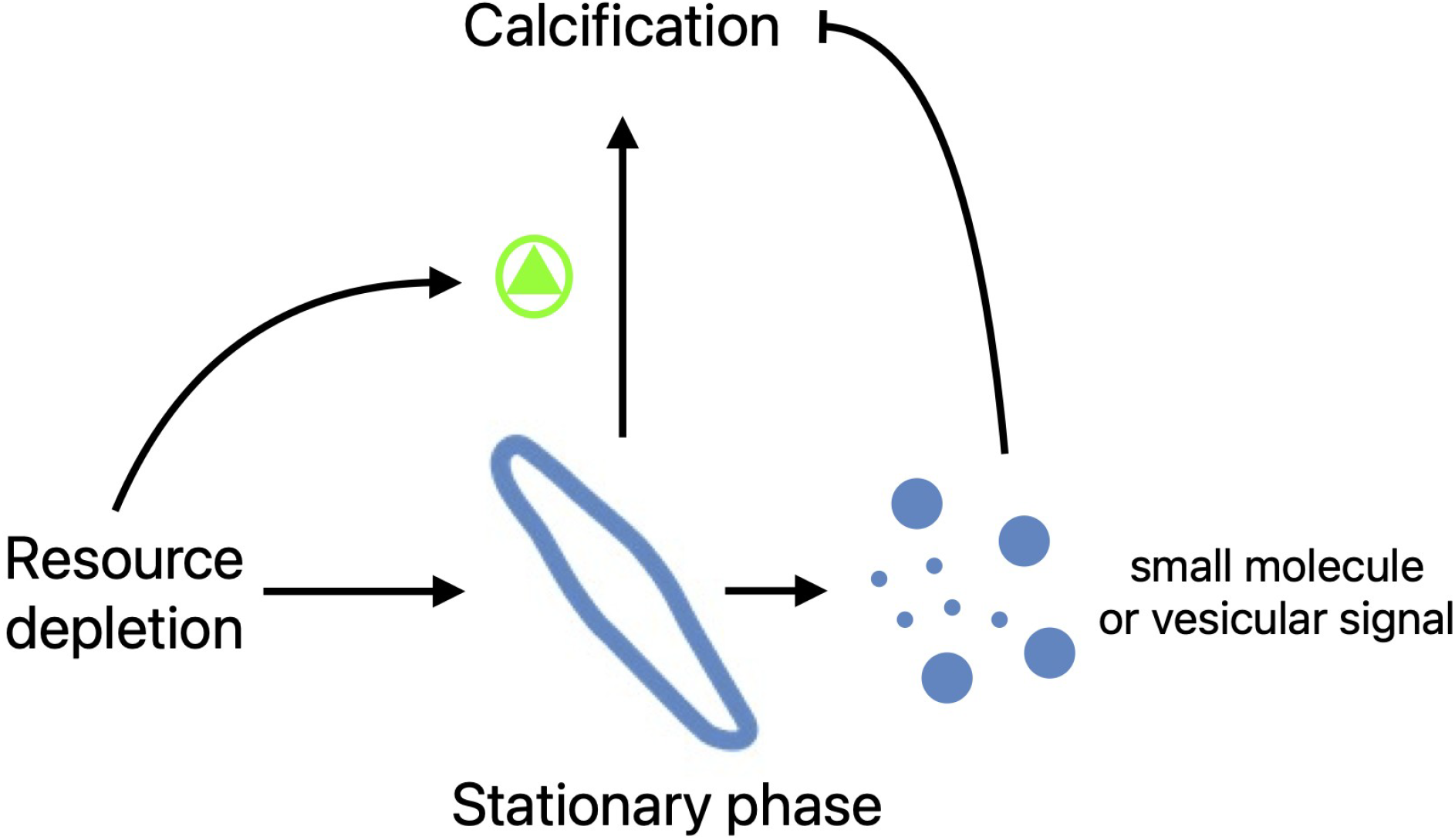
Hypothesis of the calcification process.

### Molecular mechanism of the calcification process

According to Fig. 3, the calcifying activity is one that acts in the extracellular space rather than inside the cell to be able to generate such clumps as we have observed. The micrograph also demonstrates no complex structure within the carbonate deposit, unlike coccoliths, suggesting this is a much simpler process. For the divalent carbonate precipitation to occur, the immediate local concentrations of both species of ions, the divalent Ca2+ and/or Mg2+, and the CO32-anion, would need to be actively raised above the precipitation threshold. Most photosynthetic algae are known to have some versions of carbonic anhydrase activities to promote photosynthesis, and calcium signalling is a fairly conserved function amongst eukaryotes, so there is no functional barrier to this phenomenon in vivo. Our results that autoclaved media loses the ability to generate the deposit is supportive of this hypothesis - it is known that after autoclaving, the phosphate anion and divalent cations, mostly magnesium and calcium, form insoluble precipitates and are taken out of the resources available to the culture, this result would suggest there is a minimum threshold of the divalent cations for the calcification to occur. Neither of the two possibilities, a. that the responsible enzymes or vesicles [5] are secreted and acted or reacted extracellularly, or b. the ions were directly exported by exocytosis-like mechanisms, receive more experimental support from this brief communication.

### Ecological role for the calcification activity

One can argue that the facts that Phaeodactylum are tolerant to a wide range of salinities and are mostly sampled in coastal locations [6] would suggest the genus has evolved towards the intertidal or estuarine ecology, where both salinity and exposure to air can vary wildly within a very short period of time. We propose that the newly discovered activity helps the Phaeodactylum population to secure themselves to the physical objects in their environment without being washed into the deeper part of the sea. In our hands, Phaeodactylum tricornutum cells are poor at staying in the water column when cultured without shaking, sedimenting readily. Whilst there are studies suggesting that Phaeodactylum has the activity to control cell buoyancy, travelling vertically in the water column [7-8] - some of which suggesting the environmental cue involving nutrient starvation - we postulate that this physical tethering to solid objects close to the shore line may be an alternative strategy to remain in the photosynthetically active part of seawater.

## Methods

### Cell culture media

f/2 media was prepared by dissolving 35g/L Instant Ocean to deionised water (10 MΩ), then adding the f/2 supplement (Phytoreef Ltd) according to instruction. L1* media is a simplified modification of Karas et al. [2], rather than the earliest instance where such medium was published [9]. This was prepared by dissolving 35g/L Instant Ocean to deionised water, adding the f/2 supplement as f/2, and adjusting NaNO3 and NaH2PO4 to 10x of originally published f/2 concentration [10]. Artificial Sea Water Media (ASW media) was prepared according to [11].

### Cell culturing

For tissue culture flasks, T25 for suspension cells (690195, Greiner Bio-One) were used in a PHCBi MLR-352-PE, light level 2, 16/8 light/dark cycle at constant 18°C, gentle orbital shaking. Bubbling bioreactor build: see Fig. 1a.

For raceway ponds, the ponds were cleaned with ethanol and water after each use. Media was made inside the ponds before inoculation with the paddle wheel activated to dissolve the salt components. No extra decontamination processes were made. During pond action, the ponds receive 120mmol of light at the surface of the water column, being set on a 16/8 light/dark cycle with the paddle wheel constantly on.

### Spiking and media replacement experiments

For the spiking experiment (Table 2), cultures were grown in 250mL bottle bubbling photobioreactors until they reached stationary phase, then were spiked with NaNO3 and NaH2PO4 to 10% of original concentration of fresh f/2 media. In the media replacement experiments, cultures at stationary phase were pooled, concentrated by gentle centrifugation, washed with Instant Ocean seawater, repeated centrifugation, then brought back to original volume in respective media.

### Inductively Coupled Plasma Mass Spectrometry (ICP-MS)

The analysis was performed by James Hutton Institute (Craigiebuckler, Aberdeen AB15 8QH, Scotland). 15mg samples were dissolved in 500 µL of 15% HNO_3_, then diluted with 9.5 mL of deionised water (1:20 dilution). The measurement was made on this solution.

## Notes

### Competing Interest Statement

The authors have declared no competing interest.

## References

[1] Avasthi, P., MacQuarrie, CD., Radkov, A., (2023) Inducing protoplast formation in Phaeodactylum tricornutum by silica deprivation, enzymatic treatment, or cytoskeletal inhibition, Arcadia Science (preprint)

[2] Karas, B., Diner, RE., Lefebvre SC., et al., (2015) Designer diatom episomes delivered by bacterial conjugation, Nature Comm., (6) 6925

[3] Marshall, DW., and Volcani. BE., (1971) Synchronized cultures: diatoms. Met. in enzym. (23) 85–96

[4] Murphy, E., Gorman, L.., Helliwell, K., et al., (2026) Glutamate as an extracellular signalling molecule in unicellular protists, Current Biology, (36), 9

[5] Picciotto, S., Barone, ME., Fierli, D., et al., (2021) Isolation of extracellular vesicles from microalgae: towards the production of sustainable and natural nanocarriers of bioactive compounds, Biomaterials Sci., 9 (8): 2917–2930

[6] Kräbs, G., Büchel, K., (2011) Temperature and salinity tolerances of geographically separated Phaeodactylum tricornutum Böhlin strains: maximum quantum yield of primary photochemistry, pigmentation, proline content and growth, Botanica Marina 54 231–241

[7] Erga, SR., Lie, GC., Aarø, LH., (2010) Fine scale vertical displacement of Phaeodactylum tricornutum (Bacillariophyceae) in stratified waters: Influence of halocline and day length on buoyancy control, 384, 1–2, 7-17

[8] Zhang, W., Hao, .Q., Zhu, J., et al., (2023) Nanoplanktonic diatom rapidly alters sinking velocity via regulating lipid content and composition in response to changing nutrient concentrations, Front. Mar. Sci., 10

[9] Guillard, RRL., Hargraves, PE., (1992) Stichochrysis immobilis is a diatom, not a chrysophyte, Phycologia, 32

[10] Guillard, RR., Ryther, JH., (1962) Studies of marine planktonic diatoms. I. Cyclotella nana Hustedt, and Detonula confervacea (cleve) Gran, Can. J. Microbiol. 8:229–39.

[11] Darley, WM., Volcani, BE., (1969) Role of silicon in diatom metabolism: a silicon requirement for deoxyribonucleic acid synthesis in the diatom Cylindrotheca fusiformis Reimann and Lewin.” Exp. Cell Res. 58 2-3 334–342

